# Hitching a Ride: Mechanics of Organelle Transport Through Linker-Mediated Hitchhiking

**DOI:** 10.1101/811372

**Authors:** Saurabh S. Mogre, Jenna R. Christensen, Samara L. Reck-Peterson, Elena F. Koslover

## Abstract

In contrast to the canonical picture of transport by direct attachment to motor proteins, recent evidence shows that a number of intracellular ‘cargos’ navigate the cytoplasm by hitchhiking on motor-driven ‘carrier’ organelles. We describe a quantitative model of intracellular cargo transport via hitchhiking, examining the efficiency of hitchhiking initiation as a function of geometric and mechanical parameters. We focus specifically on the parameter regime relevant to the hitchhiking motion of peroxisome organelles in fungal hyphae. Our work predicts the dependence of transport initiation rates on the distribution of cytoskeletal tracks and carrier organelles, as well as the number, length and flexibility of the linker proteins that mediate contact between the carrier and the hitchhiking cargo. Furthermore, we demonstrate that attaching organelles to microtubules can result in a substantial enhancement of the hitchhiking initiation rate in tubular geometries such as those found in fungal hyphae. This enhancement is expected to increase the overall transport rate of hitchhiking organelles, and lead to greater efficiency in organelle dispersion. Our results leverage a quantitative physical model to highlight the importance of organelle encounter dynamics in non-canonical intracellular transport.

**SIGNIFICANCE:** A variety of cellular components are transported via hitchhiking by attaching to other motile organelles. Defects in the molecular machinery responsible for organelle hitchhiking may be linked with neurodegenerative disorders. To date, no comprehensive physical models of this non-canonical mode of transport have been developed. In particular, the connection between molecular- and organelle-scale properties of hitchhiking components and their effect on cellular-scale transport has remained unclear. Here, we investigate the mechanics of hitchhiking initiation and explore organelle interactions that can modulate the efficiency of this process.

## INTRODUCTION

Regulated movement of proteins, vesicles, and organelles plays an important role in the growth, metabolism and maintenance of cellular health. These particles move within a crowded and dynamic intracellular environment, aided by a dedicated transport machinery that typically comprises molecular motor proteins walking upon a network of cytoskeletal filaments. Precise control of transport ranging over length scales from a few microns to tens of centimeters is achieved by regulating the interactions between moving and stationary cargo, motors, and other cytoskeletal structures. Defects in the regulation of organelle movement can lead to pathologies, particularly in long cells such as neurons, where axonal transport deficiencies have been implicated in neurodegenerative disorders including Alzheimer’s, amyotrophic lateral sclerosis (ALS), and multiple sclerosis(1–3).

The traditional picture of intracellular transport involves the direct attachment of cargo to adaptor proteins that recruit cytoskeletal motors, which carry the cargo processively along microtubule tracks(4–6). However, recent experimental evidence suggests that a variety of cargos such as peroxisomes, lipid droplets, messenger ribonucleoprotein (mRNP) complexes, RNA granules and the endoplasmic reticulum can attach to other motile organelles, and navigate the cytoplasm through a mode of transport known as “hitchhiking”(7–15). Hitchhiking is defined by the presence of a motor-driven “carrier” organelle which colocalizes with a cargo (the hitch-hiker) and is required for its processive transport. The ubiquity of hitchhiking cargos across systems suggests that this is a broadly applicable transport mechanism(16), whose efficiency may dictate the distribution and delivery of particles that are critical for optimal cellular function.

Previous theoretical models of canonical microtubule-based transport have focused on the distribution of cytoskeletal tracks(17), interplay between diffusive and processive transport (18, 19), characteristics of motor processivity and turning (20), and cargo behavior at microtubule intersections(21, 22). The non-canonical hitchhiking mechanism, however, is governed by fundamentally different interactions at the molecular and organelle level, as compared to classic motor-driven transport. The physical principles that underlie hitchhiking efficiency have not yet been quantitatively explored.

Although the molecular components of hitchhiking have yet to be fully identified for many cargos, linker proteins which link the hitchhiking cargo to the carrier organelle have been identified in some cases(15, 16). For example, in the filamentous fungus *Ustilago maydis*, mRNAs and their associated polysomes attach to early endosomes via an interaction between RNA-binding protein Rrm4 and early endosome-associated protein Upa1 (7–10). In both *Ustilago maydis* and *Aspergillus nidulans*, another filamentous fungus, peroxisomes hitchhike on early endosomes (12, 13). In *Aspergillus*, the protein PxdA is required for peroxisome hitchhiking(13), and is a putative linker between early endosomes and peroxisomes. In rat neurons, RNA granules hitchhike on motile lysosomes using the ALS-associated protein Annexin A11 as a linker(15). While such linker proteins have been shown to be required for hitchhiking in these circumstances, it remains unknown how their mechanical and structural properties modulate hitchhiking efficiency.

In some cell types, organelles such as peroxisomes and mitochondria have been observed to attach to microtubules when not being actively transported(23–25). Such tethering allows for regulated placement of organelles within the cell(25–28). Tethering may also enhance the ability of cargo to interact with the transport machinery, increasing the rate of initiating active runs while restricting short-range diffusion(19). Peroxisomes in particular have been found to exhibit both diffusive motion and microtubule tethering depending on cell type and context (14, 23), indicating that both modes of motion may play a role in organelle motility. Here, we explore how tethering to microtubule tracks can enhance the rate of transport initiation for hitchhiking cargos, by placing them within easy reach of passing carrier organelles.

Given the complexity of intracellular transport processes, many studies of transport have focused on the simplified geometries found in long cylindrical cellular projections(13, 14). Such projections feature polarized arrays of parallel microtubules, with few intersections and essentially one-dimensional movement of cargo. Neuronal axons and the hyphae of filamentous fungi exhibit a similar cylindrical geometry. Filamentous fungi such as *Aspergillus nidulans* are particularly amenable to genetic manipulation and imaging, providing convenient experimental systems for studies of intracellular transport (29).

In this paper we investigate the effects of cellular and cytoskeletal geometry, as well as mechanical properties of the transport machinery, on the interaction between hitchhiking cargos and carrier organelles. We develop an analytical and computational model of hitchhiking transport initiation within tubular geometries, quantifying the rate of cargo-carrier contact for a wide range of biologically feasible parameters. In particular, we focus on peroxisome hitchhiking in fungal hyphae, leveraging experimental observations to identify the relevant parameter regime. We analyze the role of linker proteins in mediating the contact between carrier and cargo organelle, and establish optimum mechanical and structural parameters for linkers that can maximize the hitchhiking initiation rate. For organelles that can tether to microtubule tracks, we quantify the potential enhancement of the hitchhiking rate due to tethering, and identify its effect on overall organelle dispersion in the cell.

## METHODS

Overdamped Brownian dynamics simulations are employed to explore the dynamics of carrier and cargo organelles as they first encounter each other for hitchhiking initiation. Our simulation framework is generally applicable to quantifying organelle encounters in a cylindrical domain, although the parameters selected here are relevant specifically for endosomes and peroxisomes in fungal hyphae. The carriers (*e*.*g*.: endosomes) are modeled as spheres of radius *r*_*e*_ = 100nm, moving in a domain of length *L* = 1*µ*m and radius *R* = 1*µ*m with periodic boundaries in the axial direction. The domain represents a section of cell around a single cargo capable of hitchhiking. In *A. nidulans* hyphae, peroxisomes are observed at an average linear density of approximately 1 organelle within a 1*µ*m long region of the hypha (see Supplemental Materials) (14), which sets the length of our simulated domain. The radius of the domain is set to match the typical radius of fungal hyphae (see Supplemental Materials for measurements in *A. nidulans*)(14).

Microtubules are modeled as *N* straight lines distributed uniformly within the domain cross section. Microtubule dynamics are not included in the current model, although they provide an interesting avenue for future study. We ignore transverse fluctuations of microtubules, given that their persistence length *in vivo* (*l*_*p*_ ≈ 30*µ*m (30)) is much longer than the domain length. The linear density of carrier organelles (*ρ*) gives the number of carriers per unit length of hypha. Our simulation includes *ρL* carriers within the simulated domain. Each carrier is attached to a microtubule track by a single zero-length stiff spring representing a molecular motor complex. The attachment point of the spring to the microtubule moves processively in either direction at a constant velocity of 2*µ*m/s, comparable to the measured velocities of fungal peroxisomes and early endosomes(13). Upon leaving the domain, the carrier organelle reappears at the other side, on a newly selected microtubule, thereby maintaining a constant carrier density while representing organelles whose typical run-length is much longer than the domain length *L*.

Cargo organelles are represented by spheres of radius *r*_*p*_ = 100nm, which either diffuse freely through the domain, or have a point on their surface attached to a fixed microtubule at the axial center of the domain. Both carriers and cargo experience Brownian forces and torques corresponding to translational diffusivity *D*_*t*_ = *k*_*b*_*T/*(6*πηr*) and rotational diffusivity *D*_*r*_ = *k*_*b*_*T/*(8*πηr*^3^), where *η* is the viscosity of the domain. The viscosity is chosen such that *D*_*t*_ ≈ 0.014*µ*m^2^*/*s for the cargo organelle, in keeping with measured diffusivities of peroxisomes in *Ustilago maydis* hyphae(14). Steric interactions between organelles and with the cylindrical boundary of the domain are implemented using a stiff harmonic potential.

The simulations are evolved forward using a fourth-order Runge-Kutta algorithm(31) with time-steps of 10^−4^s. Each simulation trial is run for a total of 5s, allowing each carrier to pass 10 times through the domain. 2500 trials are carried out for each combination of carrier density *ρ* and microtubule number *N*.

Linker proteins that mediate contact between carrier and cargo are modeled as continuous semiflexible worm-like chain (WLC) polymers (32) with varying length. Positions of the base of the linker protein are chosen uniformly on the carrier surface, and the initial linker tangent is assumed perpendicular to the surface. Using analytically calculated distributions for the end point of a WLC(33), we tabulate the spatial distribution of the probability that a cargo organelle overlaps with the tip of a linker for a given configuration of the carrier and cargo (see Supplemental Materials). Using this tabulated probability, at each time step we check whether linker-mediated contact between the carrier and cargo has occurred. This approach avoids resolving the dynamics of the linker protein configurations, working instead in the fast-equilibration limit where the position of each linker tip is sampled from its equilibrium distribution at each timestep.

For each simulation trial, we note the time until the single cargo organelle first contacts either the carrier surface or the tip of a linker protein. The empirical cumulative distribution function is used to extract an effective rate of contact. Over the simulation timescale, the cumulative distribution functions observed fit well to a double exponential form 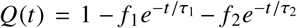 (see Supplemental Materials). The average rate of hitchhiking initiation is then defined by *k*_hit_ = (*f*_1_*τ*_1_ + *f*_2_*τ*_2_)^-1^. Variation in this initiation rate is approximated by bootstrapping(34) over all simulation trials for a given set of parameters. All error bars shown give the standard deviation in *k*_hit_ over 100 bootstrapping runs.

Brownian dynamics simulation code (in Fortran 90) and scripts for implementing linker distributions and obtaining encounter rates from simulation results are provided in a GitHub repository: https://github.com/lenafabr/hitchhiking_initiation.

## RESULTS AND DISCUSSION

### Rate of encounter with carrier organelles

The efficiency of hitchhiking transport initiation is governed in part by geometric parameters, such as the density of microtubules and carrier organelles, as well as the length and distribution of linkers on the carrier surface. In order to be picked up for a hitchhiking run, the cargo must be sufficiently close to a passing carrier to be able to engage with a linker protein. We begin first by considering the rate of encounters between a diffusing cargo organelle and a processively moving carrier. This rate corresponds to hitchhiking initiation in the limit of very short, densely packed linkers, where the entire carrier surface is capable of binding the cargo.

A Brownian dynamics simulation framework is employed to explore how the density of microtubules and carriers modulates organelle encounter in a tubular region of radius *R* = 1*µ*m with parameters relevant to peroxisome transport in hyphae (Fig. 1). The radius of the tubular region corresponds to the average radius of *A. nidulans* hyphae, as obtained from experimental measurements (see Supplemental Materials). A variable number (*N*) of parallel microtubules are uniformly scattered throughout the tubular region. A single cargo of radius *r*_*p*_ = 100nm and translational diffusivity *D*_*t*_ = 0.014*µ*m^2^*/*s represents the peroxisome and a variable linear density *ρ* of carrier organelles of radius *r*_*e*_ = 100nm move with processive velocity v = 2*µ*m/s along the microtubule tracks. Brownian forces on the carrier organelles drive fluctuations around their attachment point to the microtubules. Periodic axial boundary conditions allow for maintenance of a constant density of processively moving carriers in the local vicinity of the cargo.

**Figure 1:**
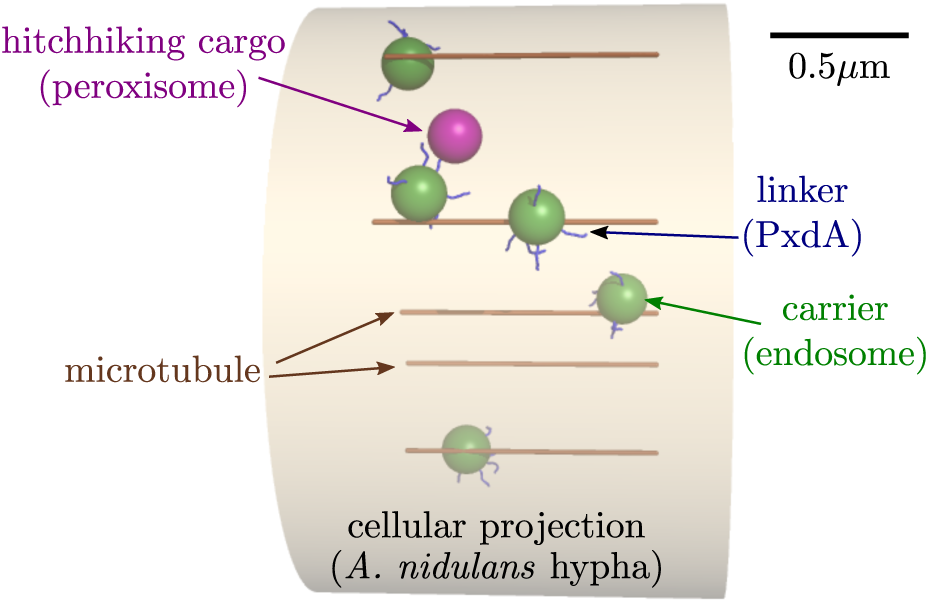
Schematic of the model for hitch-hiking initiation. A simulation snapshot is shown with all model components labeled. Specific components for peroxisome hitchhiking in *Aspergillus nidulans* are indicated in parentheses. Scale bar: 0.5*µ*m.

In order to come in contact with a carrier, the cargo must first approach sufficiently close to a microtubule track (within a distance of *r*_*p*_ + 2*r*_*e*_ = 0.3*µ*m), and then be hit by a passing carrier before moving away from the track again (Fig. 2a). For a single, centrally located microtubule, the region of proximity covers a fraction *f*_1_ = (*r*_*p*_ + 2*r*_*e*_)^2^*/*(*R*− *r*_*p*_)^2^ ≈ 0.1 of the available cross-sectional domain area. As multiple parallel microtubules are placed within the domain, their proximity regions cover an increasing fraction of the cross-sectional area. We vary the number of microtubules *N* in our simulation, randomizing the placement of each microtubule and the initial radial position of the peroxisome within the domain. Fig. 2b shows the fraction of iterations where the peroxisome starts within reach of a microtubule (equivalent to the MT-proximal area fraction *f*_*N*_), as well as *k*_MT_, an effective rate for peroxisomes initiated outside of the MT-proximal area to first reach this area (see Methods for details of rate calculations).

**Figure 2:**
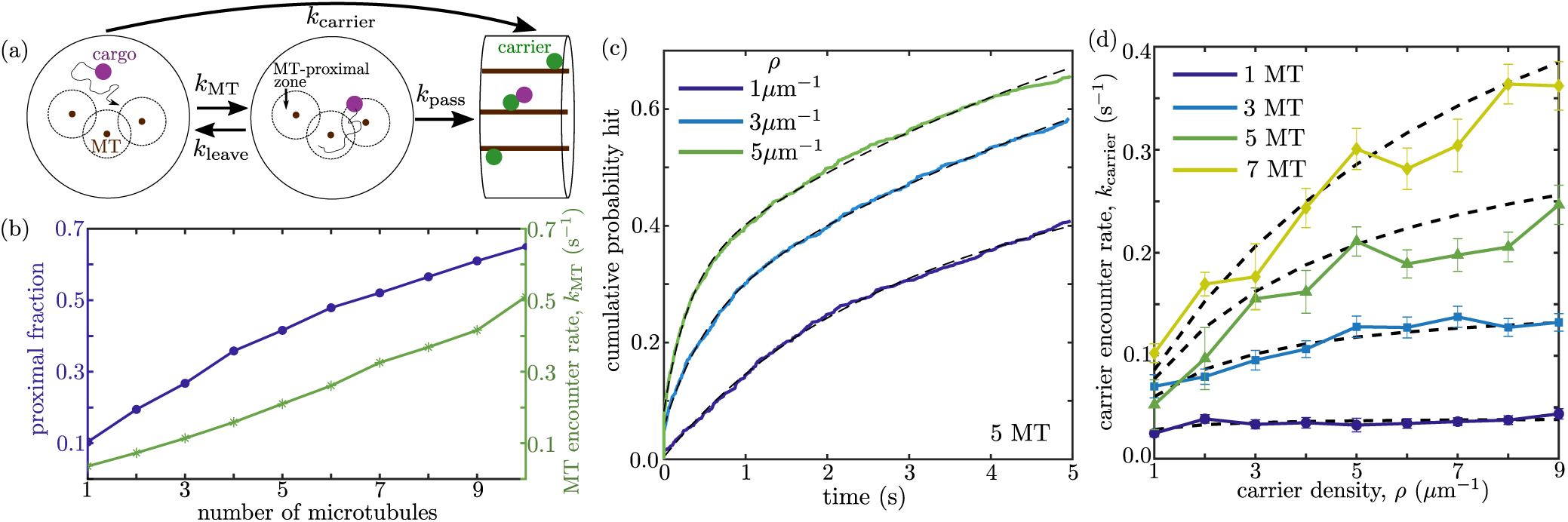
Dynamics of cargo encounter with carrier organelles. (a) Schematic model for carrier encounter, illustrating the two-step process of first entering a MT-proximal zone, then waiting for a carrier passage event. (b) Average fraction of domain cross-sectional area within distance 2*r*_*e*_ + *r*_*p*_ from a microtubule (left) and rate *k*_MT_ to encounter a microtubule if starting outside the proximal zone (right). (c) Cumulative distribution of carrier encounter times, plotted for simulations with three different carrier densities. Dashed lines give fit to a double-exponential function used to extract an effective encounter rate. (d) Average carrier encounter rate, for different numbers of microtubules and carrier density. Symbols indicate simulation results; dashed black lines show predictions from approximate kinetic model (Eq. 2).

The time for a cargo to encounter a passing carrier is governed both by the dynamics of entering and leaving the MT-proximal region (rates *k*_MT_, *k*_leave_) and the rate *k*_pass_ of carrier passage in the vicinity of a cargo that is within reach of a microtubule. Because the velocity of processive motion is rapid compared to the cargo diffusivity, we treat the carrier arrival as a constant rate process while the peroxisome is within the MT-proximal region. The rate of this arrival is given by

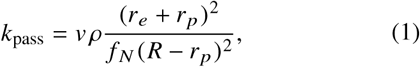

where *v ρ* is the rate at which carriers pass the axial position of the cargo and the second term corresponds to the equilibrium probability that the radial position of the cargo is within reach of the passing carrier, assuming the cargo is uniformly distributed within the MT-proximal area. The effective rate of leaving the MT-proximal area must be such that the cargo spends fraction *f* _*N*_ of its time within this area at equilibrium. Namely, *k*_leave_ = *k*_MT_[1 - *f* _*N*_]*/ f* _*N*_. These three rates allow for an approximate calculation of the waiting time for a cargo organelle distributed uniformly within the domain to first encounter a carrier, using the simplified kinetic scheme shown in Fig. 2a. The inverse of this time gives the effective carrier encounter rate:

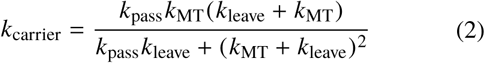

The typical time-scale for cargo-carrier encounter in simulated trajectories is obtained by fitting the computed cumulative distribution function to a double exponential process (Fig. 2c; details in Supplemental Material). As shown in Fig. 2d, effective encounter rates obtained from the simulations are well represented by the simplified kinetic model of Eq. 2.

At low carrier density, increasing the microtubule number beyond a couple of microtubules has little effect on the rate with which the cargo first encounters a carrier. In this regime, the diffusing cargo has time to enter and leave the MT-proximal region while waiting for a carrier to pass by. Each carrier passage event becomes essentially independent from the previous one, in terms of the probability that it will hit the cargo. Splitting up a fixed carrier density across more microtubules does not change the overall frequency of these independent passage events and thus has little effect on the encounter rate. By contrast, at higher carrier densities increasing the number of parallel microtubules can greatly speed up the encounter process. In this limit, carriers arrive very rapidly and the encounter is limited by the cargo approaching sufficiently close to a microtubule to enable contact. Hence, increasing *k*_MT_ by raising the number of microtubules will increase the overall encounter rate.

Similarly, when there are very few microtubules in the domain, the encounter rate is nearly independent of the carrier density. A greater frequency of carrier passage events along a single microtubule will not speed up encounter times that are dominated by the cargo coming in radial proximity of that microtubule. At higher microtubule numbers, the cargo spends most of its time within the MT-proximal region and increasing carrier density enhances the rate at which some carrier passes the cargo on a nearby microtubule.

We quantified the number of microtubule plus-ends in *Aspergillus nidulans* hyphae, and found approximately *N* ≈ 5 parallel microtubules at the hyphal tips (see Supplemental Materials). For this number of microtubules, the rate of cargo encounter with a carrier increases with the carrier linear density up to *ρ* ≈ 5*µ*m^-1^, after which it is insensitive to the presence of additional carrier organelles. We also quantified the linear density of fluorescently-tagged early endosomes in *A. nidulans* hyphae and found approximately 5 endosomes per *µ*m length of hypha (see Supplemental Materials). As approximately 55% of endosomes carry the PxdA linker protein responsible for peroxisome hitchhiking(13), we would expect the rate of peroxisome encounter with a PxdA-bearing endosome (*ρ* ≈ 2.8*µ*m^-1^) to be approximately 0.2s^-1^.

### Rate of encounter with linker proteins

Organelle hitchhiking is generally believed to involve the cargo (hitchhiker) attaching to a carrier organelle surface via one or more linker proteins (16). For the case of peroxisome transport in *A. nidulans*, the putative linker protein (PxdA) is present on a subpopulation of early endosomes and is required for peroxisome hitchhiking (13). This protein contains a long predicted coiled-coil region, which is approximately 90nm in length if fully extended(13). Given that the linker protein may be comparable in size to the organelles themselves, its length, distribution, and mechanical properties have the potential to substantially impact the efficiency of hitchhiking initiation. We thus incorporate extended linker proteins on the carrier surface into our dynamic model and proceed to explore how linker protein parameters modulate the rate at which the cargo organelle can get picked up for a hitchhiking run.

We use a multi-scale approach to integrate the linkers into our Brownian dynamics simulations. The linkers are treated as semiflexible “worm-like” chains (WLC)(32) of length *t*, with one end fixed at a given position on the endosome and the initial tangent fixed to be perpendicular to the endosome surface. It is not known whether linker proteins are capable of diffusing over the carrier surface. In the extreme case of very rapidly diffusing linkers, any close approach of the hitchhiking cargo to the moving carrier would result in an encounter with a linker, given that the proteins could explore the entire carrier surface very fast compared to the relative motion of the organelles. This limiting case approaches the situation where the entire carrier surface is capable of interacting with the cargo, as discussed in the previous section, albeit with an expanded effective carrier radius resulting from the added extension of the linker beyond the carrier surface. We focus here on the opposite limit, where a given number of linker proteins is attached to fixed points on the carrier surface with no diffusion permitted for the attachment points.

Assuming that the conformation of an individual linker protein equilibrates much faster than the large organelle movements, we employ a separation of timescales in our simulation. Specifically, we assume that each linker samples its configuration from an equilibrium distribution independently on each Brownian dynamics step. We make use of the analytically known distribution function for a WLC with fixed end orientation (33) to compute the probability that a free linker end will intersect with the hitchhiking cargo for a given position of the cargo relative to the anchoring point of the linker (Fig. 3a). These probability distributions are used to sample whether the cargo has come into contact with a linker tip during each simulation step. The hitchhiking initiation time is then taken to be the total simulation time until the first such contact with a linker tip occurs. We note that this model assumes all contacts between a linker tip and a cargo organelle lead to rapid formation of a long-lasting interaction that results in a hitchhiking run. In particular, the entire surface of the hitchhiking cargo is assumed capable of interacting with the linker protein. The hitchhiking rates discussed here are thus an upper estimate on actual initiation rates.

**Figure 3:**
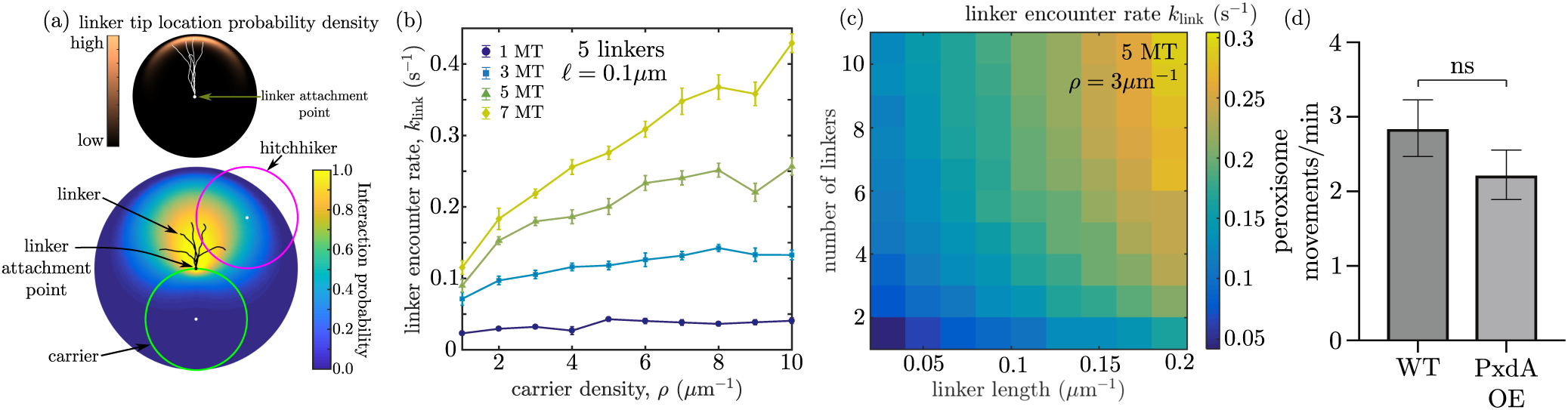
Rates of encounter with hitchhiking linkers. (a) Model of linker protein chains attached to carrier surface. Top: end distribution of a WLC of length*ℓ* = *ℓ*_*p*_, with initial end orientation fixed. Representative configurations of the linker are indicated in white. The color represents the probability density of the linker tip location. Bottom: probability of interaction between the tip of a linker (black lines) on a carrier (green) and the hitchhiking cargo (magenta). The color represents the interaction probability for each position of the hitchiker center relative to the linker attachment point. (b) Rate at which the cargo encounters the first linker tip, as a function of the carrier density and the number of microtubules. (c) Effect of linker length and linker number per carrier on the encounter rate. (d) Bar graph of the peroxisome flux in wild-type (WT) hyphae and hyphae overexpressing PxdA(Δ1-500)-TagGFP (PxdA OE). Peroxisome flux is quantified as the number of peroxisome movements across a line drawn 10*µ*m from the hyphal tip in a 1 minute movie. Wild-type hyphae had 2.84 ± 0.38 (SEM) peroxisome movements per minute, while hyphae with overexpressed PxdA(Δ1-500)-TagGFP had 2.22 ± 0.33 (SEM) peroxisome movements per minute. *p* = 0.2569, Mann-Whitney test. Error bars=SEM. *n* = 46 WT hyphae and 49 PxdA OE hyphae. See Supplementary Material for details of peroxisome flux quantification.

The rate of encounter with a linker tip shows a similar dependence on microtubule and carrier density (Fig. 3b) as the rate of coming into contact with the carrier organelle itself (Fig. 2b), discussed in the previous section. Specifically, increasing the carrier density has little effect for low microtubule numbers. Interestingly, the rates of linker contact are quite similar to the rate of carrier contact, even for a fairly small number of linkers on the carrier surface. Fig. 3c shows how the rate of linker encounter with cargo depends on the number and length of the linkers. Due to the flexibility and length of the linker proteins, only a small number of linkers (eg: ∼5, for linkers of size comparable to the putative PxdA coiled-coil region) is sufficient to obtain near maximum initiation rates. This prediction from the physical model is consistent with experimental measurements, which show that overexpressing PxdA linker proteins in *A. nidulans* does not increase the rate of hitchhiking initation (Fig. 3d).

Using the standard expansion for a nearly straight worm-like chain, the linker tip will project a typical distance Δ*z* ≈ *ℓ*[1 − *ℓ/*(6*ℓ*_*p*_)] from the carrier surface(35). For proteins of length *ℓ* = 100nm and persistence length *ℓ*_*p*_ = 100nm, a single instantaneous encounter between cargo and carrier surface is expected to yield a roughly 6% chance of any given linker on that carrier intersecting with the cargo. For 5 independent linkers, this means that approximately 30% of carrier encounters will result in immediate linker contact. This fraction is increased further because the diffusion of cargo and carrier during a passage event allows them to sample a greater fraction of each other’s surface, as has been quantified for the case of molecular diffusion towards receptor patches on a sphere(36).

In addition, by projecting beyond the surface of the carrier, the linker proteins serve as antennae, allowing contact while the cargo is further away from the carrier. This type of interaction effectively replaces *r*_*e*_ with *r*_*e*_ + Δ*z* and results in more rapid encounters. Consequently, for long and numerous linkers, the rate of initiation can be even faster than the rate of contact with the carrier. This effect arises from the extra range afforded by long linker proteins, allowing contact with the linker to occur while the cargo is still at a substantial distance from the carrier surface.

The rate at which linker contact occurs depends not only on the geometry and density of the linker proteins, but also on their flexibility (Fig. 4). In order to project beyond the surface of the carrier organelle, the linker proteins must be relatively stiff (*ℓ*_*p*_ ≳ *ℓ*). For linkers with a substantially shorter persistence length, the smaller value of Δ*z* implies that the cargo organelle would need to approach closer to the carrier surface in order to have a high likelihood of encountering the linker tip (Fig. 4b). Interestingly, when there is only one linker on each carrier, a slight optimum in linker flexibility is observed. This effect arises from the fact that the tip of a stiffer linker thermally explores a smaller area in the plane parallel to the surface of the carrier. Thus, when linker density is low, semiflexible linkers with *ℓ*_*p*_ comparable to *ℓ*_*t*_ increase the probability of an encounter with the linker tip each time the cargo approaches a carrier, above what it would be for infinitely stiff linkers. We note that the effective persistence length of coiled-coil protein structures is reported to be in the range of 30 − 170nm (37–40). While the detailed structure and mechanical properties of the PxdA linker are unknown, this protein appears to fall in the expected range for an efficient hitchhiking linker – namely, it has a predicted coiled-coil domain with comparable persistence length and end-to-end length. The coiled-coil domain of PxdA is thus expected to have sufficient stiffness that would allow the protein to project beyond the endosome surface, while remaining sufficiently flexible so that the linker tip could explore a substantial area around its attachment point.

**Figure 4:**
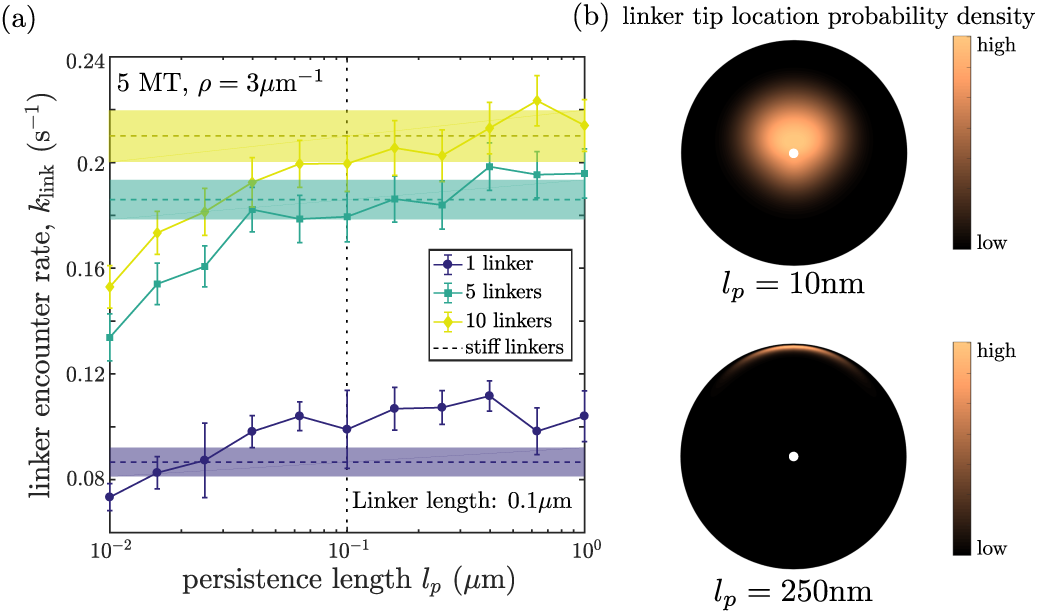
Dependence of encounter rate on linker flexibility. (a) Rate at which cargo encounters a linker tip, as a function of linker persistence length for different linker numbers. Dashed lines with shaded areas denote contact rate for infinitely stiff linkers along with standard error. Dotted vertical line denotes *l*_*p*_ = 0.1*µ*m, used in all other simulations. (b) End distribution of a WLC of length *ℓ* = 100*nm* with initial orientation fixed, highlighting the increased extension Δ*z* yet smaller area coverage of stiffer linkers. Top: *l*_*p*_ = 10nm, bottom: *l*_*p*_ = 250nm

### Tethering to microtubules enhances the rate of hitchhiking initiation

A number of organelles, including mitochondria, melanophores, and peroxisomes are known to become tethered to microtubule tracks by regulatory proteins that are crucial for maintaining their cellular distribution(23, 25, 28, 41, 42). Such tethering not only helps localize organelles along extended cell regions (as in neurons) (27, 41) but is also thought to facilitate interactions between multiple organelles by restricting their three-dimensional diffusion through the cytoplasm(28). In the case of hitchhiking cargos such as peroxisomes in *Aspergillus* and *Ustilago*, tethering to a microtubule has the potential to enhance the rate of hitchhiking initiation by eliminating the time spent out of reach of the microtubule-bound carrier organelles. We explore the effect of cargo tethering in the context of our hitchhiking initiation model by attaching the cargo surface to a randomly selected microtubule and quantifying the rate at which the cargo first encounters the carrier surface or a carrier-borne linker protein.

Tethering of the cargo to a microtubule substantially increases the rate of encounter with a carrier organelle, particularly in the case of low microtubule density (Fig. 5a). This enhancement arises from two related effects. First, the lower volume available to the cargo makes it much more likely in equilibrium (and hence at the start of the simulation) that the cargo starts in close contact with a carrier organelle. This is particularly the case for high carrier densities and low microtubule numbers, where the carriers cover nearly the entire volume available to tethered cargo. A related effect is that, even for cargos that start far from any carrier, the need to first reach a microtubule track is eliminated (*k*_MT_ → ∞), so that the rate of carrier encounter becomes comparable with the rate of carriers passing a hyphal cross-section along the same (or nearby) microtubule.

**Figure 5:**
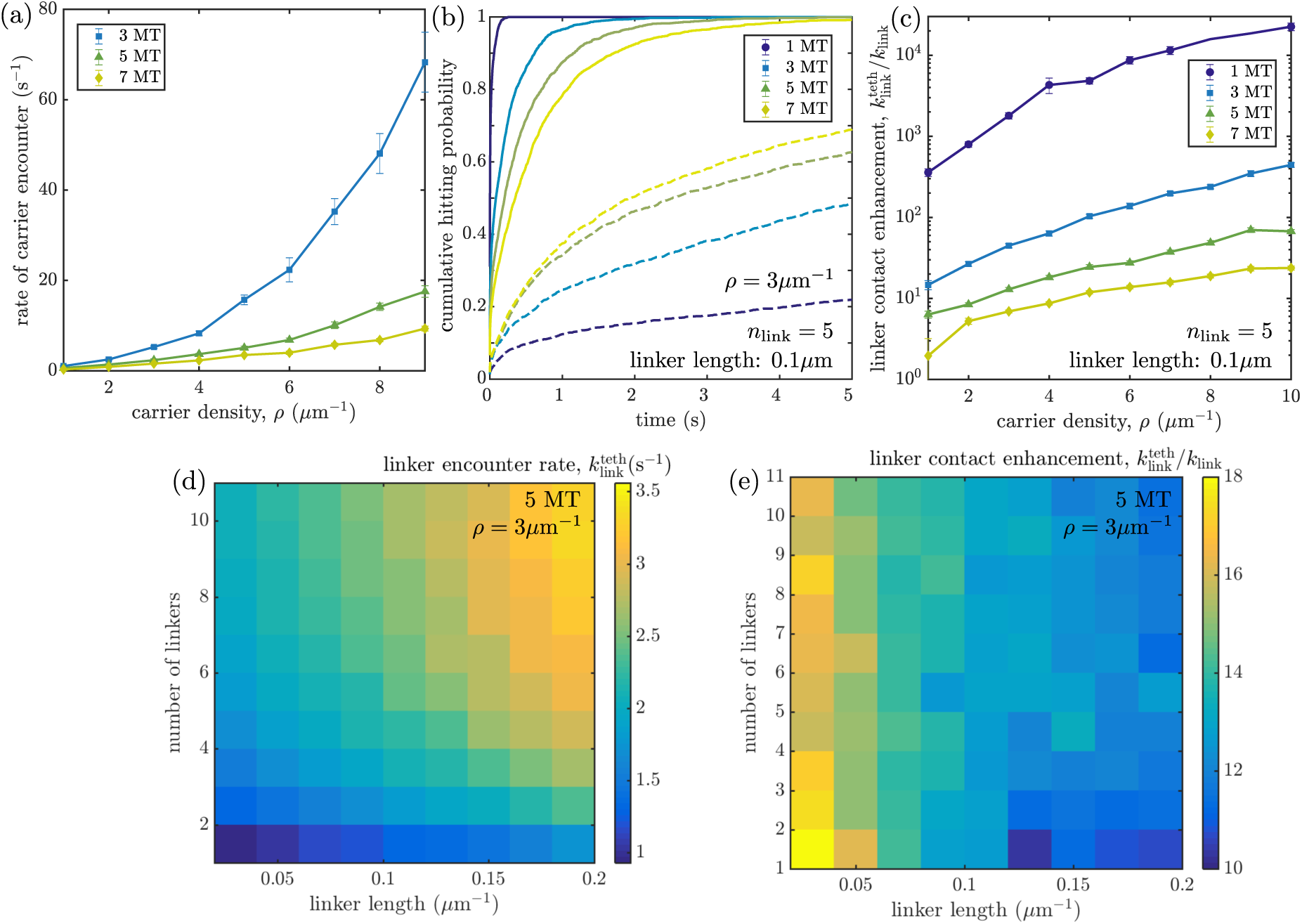
Effect of cargo tethering to microtubule on encounter rates with carrier organelles and their linker proteins. (a) Rate of encounter with a carrier organelle, for different numbers of microtubules in the domain. (b) Cumulative hitting probability with the tip of a linker protein, for tethered (solid lines) and untethered cargo (dashed lines), showing more rapid encounters in the tethered case. (c) Enhancement of overall contact rate with a linker protein tip, due to tethering of cargo. (d) Encounter rate between a tethered cargo and linker protein tip, as a function of linker length and number. (e) Ratio of encounter rates with linker protein tips, for a tethered versus diffusive cargo.

When the hitchhiking cargo is tethered to a microtubule, increasing the number of microtubules along the hypha greatly slows the rate of encounter with a passing carrier. Because we assume a constant linear density of carrier organelles per length of hypha, additional microtubules provide more options for where the carriers are located in the hyphal cross-section, diverting some of them away from the one microtubule to which the cargo organelle is attached. Hence, fewer microtubules makes it more likely that each carrier will come in contact with the cargo as it passes the relevant cross-section of the hypha.

When hitchhiking initiation requires encounter with the tip of a linker protein, tethering of the cargo can also greatly increase the rate at which such encounters occur (Fig. 5b-e). Unsurprisingly, tethering is most beneficial at low microtubule numbers and short linker lengths, where facilitating the rate at which the cargo comes near the carrier surface has a large effect on the hitchhiking initiation. For high microtubule numbers and long linkers, the effect of tethering is less pronounced because even freely diffusing cargos spend most of their time within the maximum distance (*r*_*p*_ + 2*r*_*e*_ + Δ*z*) of the microtubule tracks that allows for hitchhiking initiation during carrier passage events. As seen in Fig. 5a,c, tethering plays a greater role when the carrier density is high, since it is in this regime where contact with diffusive cargo is rate-limited by the cargo coming in proximity of a microtubule track. At lower carrier densities, the encounter time is dominated by waiting for a carrier passage event and tethering to a microtubule has less effect.

For the typical microtubule and PxdA bearing early endosome density observed in *A. nidulans* hyphae (*N* ≈ 5, *ρ* ≈ 3*µ*m^-1^; see Supplemental Materials), and for the predicted PxdA coiled-coil linker length (*ℓ* ≈ 90nm), tethering of peroxisomes is expected to increase the rate of hitchhiking initiation by more than 12-fold (Fig. 5e), even for very few linkers present on the endosome surface. Peroxisome hitchhiking in hyphae can thus be greatly enhanced by attaching the peroxisomes to microtubules. The enhancement remains substantial even if the peroxisomes are assumed to be much larger (8-fold enhancement for *r*_*p*_ = 300nm; see Supplemental Material). Published kymographs of labeled peroxisomes in pxdA-knockout hyphae(13) hint that peroxisomes exhibit little axial motion over time periods of up to 10 sec. While time-sampling limitations of this data preclude a definitive demonstration of tethering, the observed motion is not inconsistent with these organelles being attached to stationary cellular structures. Furthermore, in human cells, the peroxisomal membrane protein PEX14 has been shown to bind to tubulin and to be critical for peroxisome motility along microtubules(23). There is thus reason to propose that fungal peroxisomes may also be attached to microtubules and, as shown here, that this tethering may contribute to their ability to hitchhike throughout the hypha.

We note that for parameters relevant to peroxisomes in *A. nidulans* hyphae, the expected rate of encounter with PxdA linkers on an endosome is 2 − 3s^-1^, depending on the density of the linker proteins. This rate is well above the initiation rate of 0.02s^-1^ estimated from the small fraction (∼ 5%) of peroxisomes in *A. nidulans* hyphae engaged in hitchhiking runs(13, 19). Even without tethering, the approximate linker encounter rate of 0.2s^-1^ is an order of magnitude higher than the estimated rate of hitchhiking initiation. This discrepancy implies that only a small fraction (1 − 10%) of encounter events with the PxdA linkers result in the successful initiation of a hitchhiking run. Such a low success rate may arise from a variety of biological or mechanical reasons. In particular, it is unclear if PxdA is the actual linker between peroxisomes and early endosomes, or if it is only one component of a hitchhiking apparatus. If PxdA is the primary linker protein, it is unclear if the coiled-coil region is consistently in a fully extended form or if it takes on multiple conformations that may make it less capable of readily interacting with hitchhiking cargo. Other proteins may also regulate PxdA activity or be involved in linking peroxisomes to early endosomes in other ways. For example, only a certain percentage of peroxisomes may display a binding receptor for PxdA, creating a smaller pool of peroxisomes competent to hitchhike. As the proteins composing the hitchhiking apparatus are unknown, the strengths of contacts between the peroxisome and early endosome are unclear and may be weak. Finally, if peroxisomes are indeed tethered to microtubules, there may be a high energy requirement to break those tethers and initiate a hitchhiking event.

Given all of these potential effects beyond the encounter of a hitchhiking cargo with a linker protein, our calculation yields only an upper bound on the rate of hitchhiking initiation. We would expect this maximum rate to be modified by a prefactor that describes the probability of successful hitchhiking initiation on each given encounter, incorporating the details of linker and tether interactions. The geometric and mechanical parameters described here determine the frequency of opportunities for hitchhiking to occur. The reported initiation rates and enhancement from tethering apply to the limiting case of easily broken tethers and linker proteins that can bind to the entire peroxisome surface.

### Effect of initiation rate on overall cargo dispersion

Our exploration of the dynamics of cargo and carrier encounters indicates that tethering of cargo to microtubules can greatly enhance the rate of hitchhiking initiation. However, tethering of the cargo also inhibits its ability to move diffusively throughout the domain. Therefore, we sought to determine how these two competing factors (hindered diffusion but increased hitchhiking rate due to tethering of a cargo) would balance each other to affect the long-range dispersal of hitchhiking cargo organelles throughout a cylindrical region. To address this, we switch to a simplified, analytically tractable model which focuses on the motion of hitchhiking cargo organelles along the hyphal axis. Specifically, we leverage the one-dimensional “halting creeper” model(19) where cargo organelles are treated as point particles subject to multimodal transport. The particles exhibit memoryless stochastic switching between bidirectional processive motions (with velocity v, starting rate *k*_start_, and stopping rate *k*_stop_), interspersed with diffusive periods of diffusivity *D* (Fig. 6 inset). The more detailed mechanical model for hitchhiking initiation described in previous sections allows us to calculate the effective starting rate *k*_start_ with and without tethering of the cargo organelles to microtubules.

**Figure 6:**
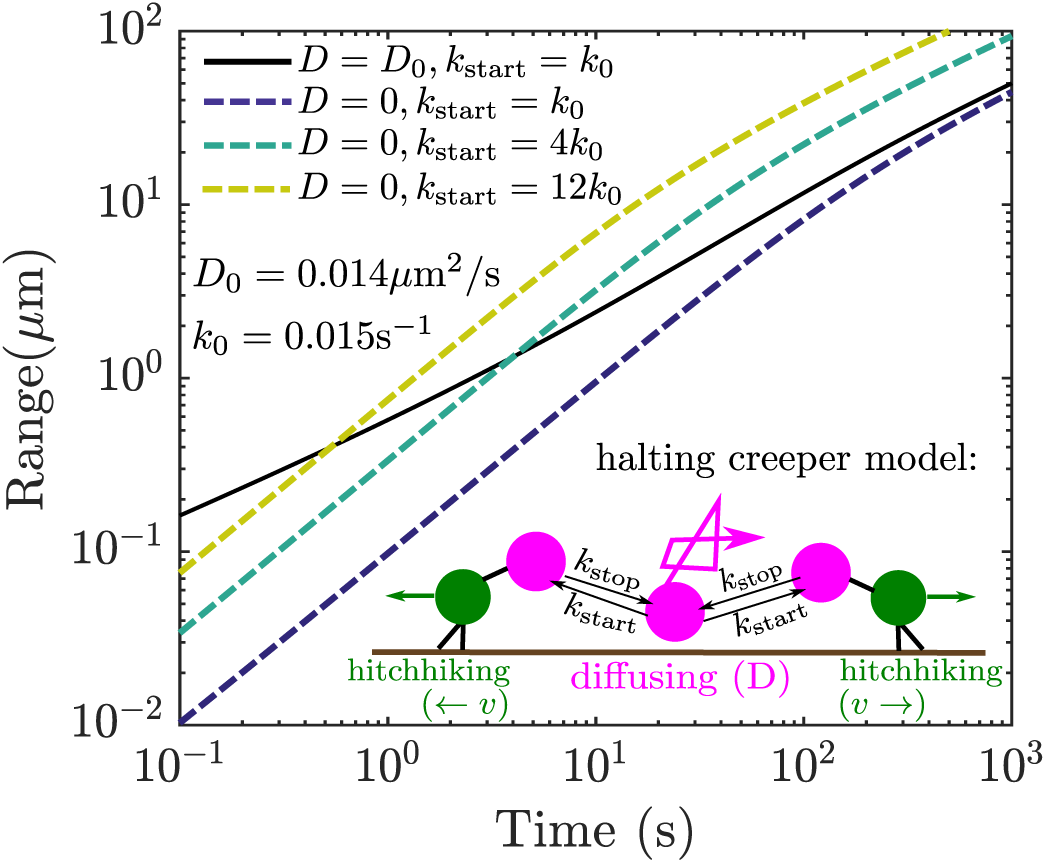
Range of a halting creeper particle engaged in bidirectional runs alternating with diffusive or tethered periods. Solid black line gives range for a particle that diffuses freely with diffusivity *D*_0_, between processive runs initiated with a rate *k*_0_. Dashed lines give range for particles that are tethered when not engaged in an active run, but with increasing initiation rates.

In prior work(19), we developed an analytical expression for the range of an exploring “halting creeper” particle. Particle range constitutes a metric of interest for intracellular transport processes because it enables direct calculation of the average time for any target in the cell to be found by the first of a uniformly distributed set of particles. It can be shown(19) that the range of a halting creeper transitions between a ballistic regime [range *Z* (*t*) = *fvt*] and an effectively diffusive regime 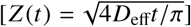, where *D*_eff_ = (1 - *f*) *D* + *fv*^2^*/k*_stop_ is an effective long-time diffusivity, and *f* = *k*_start_*/*(*k*_start_ + *k*_stop_) is the equilibrium fraction of time in processive motion. The transition to the long-range diffusive regime occurs at a time *t*\*\* = 16*D*_eff_*/*(*π f* ^2^v^2^). Increasing the starting rate for processive motion (*k*_start_) enhances the overall particle range at long times (above *t*\*\*), while decreasing the transition time where the effectively diffusive motion sets in. By contrast, in the absence of tethering, short-time motion is enhanced, creating a separate regime dominated by diffusion only with range 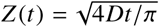.

In Fig. 6 we show the expected range of particles with different combinations of diffusivity and processive starting rate. The solid line represents approximate parameters relevant to the motion of peroxisomes as measured in *Ustilago maydis* fungal hyphae, where the peroxisomes appear to exhibit diffusive, untethered motion between processive runs(14). Namely, we set diffusivity *D*_0_ = 0.014*µ*m^2^*/*s, hitch-hiking velocity v = 1.9*µ*m/s, and run-length v *k*_stop_ = 6.5*µ*m. The initiation rate for processive runs is estimated at *k*_0_ = 0.015s^-1^, such that approximately 5% of peroxisomes are expected to be hitchhiking at any given time(14). Tethering of such particles to a microtubule will reduce the diffusivity to zero, but can enhance the starting rate by a factor of 12-fold, for the case of 5 parallel microtubules (see Fig. 5e). We thus plot how such increased processive starting rate due to tethering can enhance the range of spreading particles over time. We note that even a 4-fold increase in the starting rate raises the range of the particles above a time-scale of a couple of seconds. The 12-fold increase estimated from our hitch-hiking simulations is expected to raise overall long-time particle range by a factor of about 3-fold.

The halting creeper model thus provides insight into the relation between organelle dispersion, the rate of hitchhiking initiation, and its enhancement due to tethering. It also highlights the possible consequences of a breakdown of the tethering mechanism leading to inefficient dispersion of organelles.

## CONCLUSIONS

In this work, we describe a computational framework for delineating the key physical parameters that govern the efficiency of hitchhiking initiation. Using an analytical approach, we delineate the effects of the geometry of the transport machinery on the rate of encounter between a carrier and a hitchhiking organelle. In particular, we focus on the effects of the number of cytoskeletal tracks upon which the carrier organelles move and the linear density of the carriers. We show that encounter rates are nearly independent of the carrier density for low microtubule numbers, where the process is dominated by the ability of the diffusing cargo to come within proximity of a microtubule track. Splitting up the same carrier density across larger number of microtubules can improve the encounter rate by increasing the cross sectional coverage by the moving carriers.

In some cells, linker proteins are known to mediate the contact between a carrier and a hitchhiking organelle. We calculate the rate of encounter between a carrier and a hitchhiking organelle as a function of the length and the number of linkers on the carrier. Our results show that very few linkers of moderate length are sufficient to saturate the contact rate. This result helps explain experimental measurements showing that overexpression of PxdA linker protein does not increase the hitchhiking frequency of peroxisomes in *A. nidulans* fungal hyphae. Further exploring the effects of linker flexibility, we show that moderately stiff linkers provide optimal contact rates between the carrier and hitchhiking cargo, by allowing the linker tips to explore large volumes of space while extending substantially above the surface of the carrier.

Leveraging our simulation framework, we study the effect of tethering organelles to microtubules on the initiation of hitchhiking. The increased proximity to moving carriers results in a large enhancement of the contact rate, an effect that is particularly pronounced for small microtubule numbers and high carrier densities. Based on this enhancement in the initiation rate, we compute the increased range covered by organelles exploring the cell through rare, sporadic hitchhiking runs. Our results show that tethering can substantially increase the amount of intracellular space explored over time-scales of seconds or higher, despite restricting diffusive transport.

Our computational framework is generally applicable to any transport process that relies on attaching to a carrier organelle, either directly or through stiff or flexible linker proteins. While we focus on the simple geometry of a cylindrical domain, the parameters employed here (carrier density, density of parallel microtubules) are local in nature. Hence, the initiation rates found can be applied to any system where microtubules are arranged in a parallel fashion around the current position of the cargo organelle. This includes cellular projections such as fungal hyphae and neuronal axons and dendrites, as well as micron-sized regions of the cell soma with no microtubule intersections. Hitchhiking initiation rates in the vicinity of intersecting microtubules are left as an extension of interest for further study.

## Supporting information

Supplemental information regarding details of experimental and computational methods.

## AUTHOR CONTRIBUTIONS

SSM and EFK conceived and designed the research, and developed the model. SSM performed imaging studies, analyzed imaging data, and implemented simulations. JRC and SLRP generated experimental data and performed imaging studies. All authors contributed to data interpretation and writing of the manuscript.

## ACKNOWLEDGMENTS

We thank Cassandra Niman and Hiroyuki Hakozaki for assistance with lattice light sheet imaging and processing as well as the Nikon Imaging Center at UC San Diego for help with imaging and data analysis.

SSM acknowledges funding via a predoctoral fellowship from the Virtual Molecular Cell Consortium / Center for Transscale Structural Biology and Biophysics; EFK was supported by a fellowship from the Alfred P. Sloan Foundation and funding from the National Science Foundation CAREER Award Program (#1848057). JRC is funded by a postdoctoral fellowship from the National Institutes of Health (F32GM126692) and SLRP is funded by the Howard Hughes Medical Institute and the National Institutes of Health (R01GM121772).

## SUPPLEMENTARY MATERIAL

An online supplement to this article can be found by visiting BJ Online at http://www.biophysj.org.

## SUPPORTING CITATIONS

References (43–53) appear in the Supplementary Material.

