## Supplemental information regarding details of experimental and computational methods. for "Hitching a Ride: Mechanics of Organelle Transport Through Linker-Mediated Hitchhiking"

### Supplementary Material for
#### Hitching a Ride: Mechanics of Organelle Transport Through Linker-Mediated Hitchhiking

S. S. Mogre, J. R. Christensen, S. L. Reck-Peterson, E. F. Koslover

#### S1. *ASPERGILLUS NIDULANS* GROWTH CONDITIONS

*A. nidulans* strains were grown on yeast extract and glucose media agar gum plates for maintenance [1]. *A. nidulans* spores were then transferred prior to imaging to 1% glucose minimal media [2] supplemented with 1 mg/ml uracil, 2.4 mg/ml uridine, 2.5 $\mu$g/ml riboflavin, 1 $\mu$g/ml para-aminobenzoic acid, and/or 0.5 $\mu$g/ml pyridoxine if required.

For TIRF microscopy of mature hyphae overexpressing PxdA($\Delta$1-500)-TagGFP from the AlcA promoter, spores were inoculated onto minimal media agar gum plates with 100 mM threonine (without glucose) for 22-25 hours at 37°C. Colonies were excised from agar plates and inverted onto an eight-chambered Nunc Lab-Tek II coverglass (Thermo Fisher Scientific) for imaging. PxdA($\Delta$1-500)-TagGFP expression was confirmed by observation of motile puncta in the 488-channel (imaged as described in Section S3 c)).

For lattice light sheet and spinning disk microscopy of *A. nidulans* germlings, *A. nidulans* spores were resuspended in 1 mL of 0.01% Tween-80. The spore/Tween-80 solution was then added 1:1000 to 1% glucose minimal media supplemented with 1 $\mu$g/ml para-aminobenzoic acid on 1.5 thickness 5mm circular coverglass, and incubated for 20-24 hours at 30°C.

#### S2. *ASPERGILLUS NIDULANS* STRAIN CONSTRUCTION

*Aspergillus nidulans* strains used in this study are listed in Table I. Strain RPA402 expressing EbA-mCherry and TubA-GFP was created through genetic crossing, as previously described [4]. Strain RPA1205 overexpressing PxdA($\Delta$1-500)-TagGFP was created by homologous recombination of a single copy at the *wA* (white) locus using *AfpyrG* (*Aspergillus fumigatus pyrG*) selection into a strain lacking the *A. nidulans* homolog of human KU70, *nkuA* [2]. A PxdA construct without the first 500 amino acids was chosen and created following difficulties with cloning a GFP-tagged

| Strain | Genotype | Source |
|---|---|---|
| RPA288 | yA::[gpdA(p)-mcherry-FLAG-Pts1::Afpyro], pyroA4, pyrG89, nkuA::Bar | Salogiannis et al. 2016 [3] |
| RPA402 | wA, [EbA-mCherry-Afribo], [TubA-GFP-Afpyro], pyroA4, pyrG89, pabaA1, nkuA::argB | This study |
| RPA495 | yA::[gpdA(p)-mcherry-FLAG-Pts1::Afpyro], [TagGFP2::rabA:: AfpyrG], pyroA4, pyrG89, pabaA1, nkuA::argB+ | Salogiannis et al. 2016 [3] |
| RPA1205 | yA::[gpdA(p)-mcherry-FLAG-Pts1::Afpyro], pyroA4, wA::[AlcA(p)-pxdA($\Delta$1-500)-tagGFP::pyrG], pyrG89, nkuA::Bar | This study |

Table I: *A. nidulans* strains used in this study

PxdA full-length construct, and because PxdA without the first 500 amino acids rescues a *pxdA*Δ strain similarly to full-length PxdA (unpublished). The PxdA(Δ1-500)-TagGFP DNA construct contains an AlcA promoter sequence [5] followed by the PxdA gene without the region encoding the first 500 amino acids [3], TagGFP codon optimized for *A. nidulans*, *pxdA*'s native 3′ UTR, and the *AfpyrG* selectable marker, all flanked by 1 kb *wA* locus arms of homology. These fragments were all inserted into the Blue Heron Biotechnology pUC vector at 5′ EcoRI and 3′ HindIII restriction sites using isothermal assembly [6]. The plasmid was confirmed by sequencing, and linear DNA to be transformed into *A. nidulans* was created by PCR using primers at the 5′ and 3′ ends of the *wA* arms of homology.

#### S3. IMAGING METHODS

##### a. Lattice light sheet microscopy

*A. nidulans* germlings are grown on 1.5 thickness 5mm circular coverglass as described in Section S1, then mounted and imaged on a home-built lattice light sheet microscope to obtain timelapse z-stacks of endosomes and peroxisomes. Our home-built lattice light sheet microscope was constructed following the design described by Chen et al. [7] and detailed design information provided by the Betzig group at the Howard Hughes Medical Institute Janelia Research Campus. The optical layout was modified yet retained the relative optical component locations and optical performance of the original layout. A square lattice pattern, corresponding to 73 Bessel beams, was displayed on a SLM (Forth Dimension Displays, SXGA3DM). A 488nm laser (Coherent Genesis MX) with 0.2mW total output was shaped with two cylindrical lens pairs to illuminate the pattern. The Fourier transform of the resulting beams was projected by a 500mm lens onto an annular mask, conjugate to the back focal plane (BFP) of the excitation objective lens. The annular mask used, corresponding to numerical aperture 0.55(outer diameter) and 0.44(inner diameter) of the excitation objective lens, spatially filtered the pattern to remove unwanted diffraction orders. This annular mask provided for a 10$\mu$m beam waist (FWHM) and sheet thickness at the center of 0.92$\mu$m, measured by scanning and imaging 0.2$\mu$m beads (Thermo Fisher Scientific F8811). The BFP was projected onto galvo mirrors and used to dither the lattice pattern in the x direction continuously over a 30$\mu$m range in 0.15$\mu$m step for even sheet illumination. The fluorescence signal was imaged with a sCMOS Camera (Hamamatsu Photonics Orca Flass4.0 v3) through a bandpass filter ET525/50m (Chroma Technology) with a 20msec exposure time. All imaging was performed at room temperature. To acquire 3D images the sample was moved in 0.98$\mu$m steps over a 19.6$\mu$m range by the piezo stage it was mounted on. This corresponds to 0.51$\mu$m steps and a 10.2$\mu$m range in the detection optical axis because of a 31.5° angle between the stage axis and the light-sheet plane. Following data acquisition, the image volumes underwent rotation and deconvolution based on a measure PSF with a custom cudaDeconv software (`https://github.com/dmilkie/cudaDecon`) using the Richardson-Lucy deconvolution algorithm distributed by the Betzig group and LLSpy software developed by Dr. Tally Lambert at Harvard University (`https://github.com/tlambert03/LLSpy`) to automate the process. Maximum intensity projections of the deconvolved and rotated data were created using FIJI/ImageJ [8, 9].

##### b. Spinning disk microscopy

*A. nidulans* germlings were imaged using a Yokogawa W1 confocal scanhead mounted to a Nikon Ti2 microscope with an Apo TIRF 100x 1.49 NA objective. The scope was run with NIS Elements using the 488nm and 561nm lines of a six-line (405nm, 445nm, 488nm, 515nm, 561nm, and 640nm) LUN-F-XL laser engine and a Prime95B camera (Photometrics). Image channels were acquired sequentially using bandpass filters for each channel (525/50 and 595/50). Z-stacks were acquired using a piezo Z stage (Mad City Labs). The z-range used to image a field of germlings was set manually depending on germling extension from the coverglass surface.

##### c. TIRF microscopy

For TIRF imaging, time-lapse images were collected using a TIRF 100x /1.49 oil immersion objective on an inverted epifluorescence Ti-E microscope with Perfect Focus system (Nikon) and controlled by NIS-Elements software (Nikon). Stage position is controlled by a ProScan linear motor stage controller (Prior). GFP or mCherry fluorescence was excited by a 488-nm (50 mW) or 561-nm (50 mW) laser line, respectively. Excitation and emission paths were filtered with the appropriate single bandpass filter cubes (Chroma) and emitted signal was detected using an EM-CCD camera (Andor, iXon Ultra 888).

#### S4. ESTIMATING PARAMETERS FOR *A. NIDULANS* HYPHAE

##### a. Estimating linear density of peroxisomes and endosomes

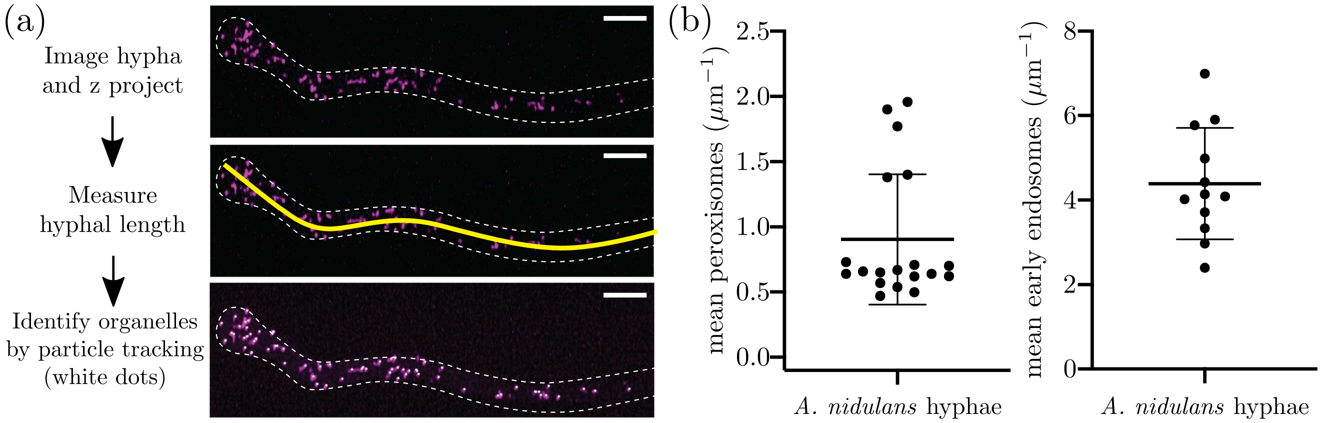


Figure S1: (a) Schematic of organelle density quantification. (Top) Maximum intensity z-projection of an *A. nidulans* germling expressing mCherry-Pts1 (peroxisome marker). White dotted line shows outline of the hypha. (Middle) Yellow line shows example hyphal length measurement. (Bottom) White dots represent each peroxisome identified . Scale bar, 5$\mu$m, (b) Scatter plots of organelle density per $\mu$m hyphal length. Each circle represents the organelle density of one hypha quantified in the 10th frame of a timelapse movie. Mean peroxisome density was $0.90 \pm 0.11$ (SEM) per $\mu$m hyphal length. Mean early endosome density was $4.57 \pm 0.41$ (SEM) per $\mu$m hyphal length. Error bars=SD. $n = 19$ hyphae for peroxisome density measurements and 12 hyphae for early endosome density measurements.

To calculate endosome or peroxisome density, lattice light sheet images of an *A. nidulans* strain expressing TagGFP-RabA (endosomes) and mCherry-Pts1 (peroxisomes) were deconvolved and z-aligned as described in Section S3 a. Maximum intensity projections were then created using FIJI/ImageJ[8]. As our simulations take into account organelle (early endosome or peroxisome) density per unit hyphal length, the organelle density was quantified by measuring the length of the hypha and quantifying the number of early endosomes or peroxisomes within that hyphal length. The length of the hyphae within the field of view was measured by manually drawing a line along the hyphal length using FIJI/ImageJ (Fig. S1) and taking a measurement. Early endosomes or peroxisomes were identified within the same regions using the Crocker-Grier 3D particle tracking algorithm [10], implemented via publicly available code[11]. The number of organelles per unit length was then calculated to determine organelle density.

##### b. Estimating microtubule number

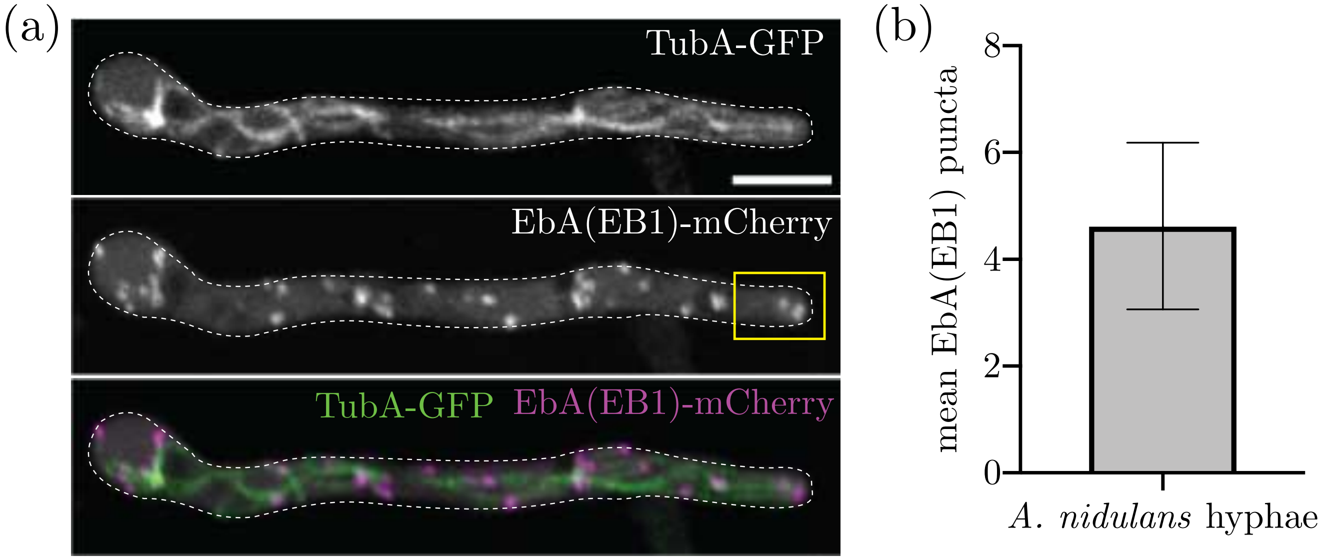


Figure S2: (a) Maximum intensity z-projection of an *A. nidulans* germling expressing TubA-GFP (microtubules) (top) and EbA(EB1)-mCherry (microtubule plus-end marker) (middle) and a merged image (bottom). White dotted line denotes outline of *A. nidulans* germling. Yellow box denotes region in which EbA(EB1) puncta were counted (the region between the hyphal tip and the nuclei nearest the hyphal tip). Scale bar, 5$\mu$m. (b) Bar graph of number of EbA(EB1) puncta in the region between the hyphal tip and the nucleus nearest the hyphal tip. Mean number of EbA(EB1) puncta was $4.62 \pm 0.14$ (SEM). Error bars=SD. $n = 133$ hyphae.

To quantify microtubule number, maximum intensity projections of spinning disk images from *A. nidulans* hyphae expressing EbA-mCherry (microtubule plus-end marker) and TubA-GFP (microtubules) were created using FIJI/ImageJ[8]. Microtubule number was quantified by manually counting EbA tips within the region between the nucleus nearest the hyphal tip and the hyphal tip. The nucleus was identified as a dark circular region in the EbA background fluorescence. If the first visible nucleus closest to the hyphal tip was further than 10 $\mu$m from the hyphal tip, then the EbA spots within 10 $\mu$m of the hyphal tip were counted.

##### c. Estimating the diameter of *Aspergillus nidulans* hyphae

To quantify the diameter of *A. nidulans* hyphae, lattice light sheet images of an *A. nidulans* strain expressing TagGFP-RabA (endosomes) and mCherry-Pts1 (peroxisomes) were deconvolved and z-aligned as described in Section S3 a. Five measurements of hyphal diameter were taken along each hypha by manually drawing a line across the hypha width and measuring using FIJI/ImageJ.

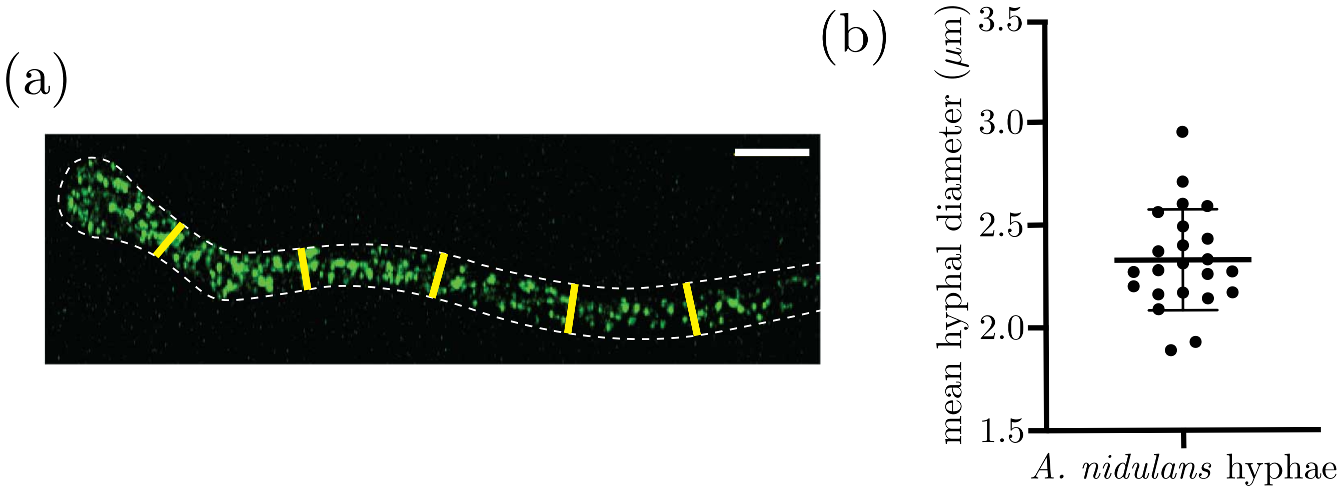


Figure S3: (a) Schematic of hyphal diameter measurements. Maximum intensity z-projection of an *A. nidulans* germling expressing TagGFP-RabA (early endosomes). White dotted line shows outline of the hypha. Five diameter measurements were taken along each hypha, represented by yellow lines. Scale bar, 5$\mu$m. (b) Scatter plot of *A. nidulans* hyphal diameter measurements. Each circle represents the average of five measurements for one hypha. Mean hyphal diameter was $2.33 \pm 0.25$ (SEM) $\mu$m. Error bars=SD. $n = 23$ hyphae.

##### d. Estimating number of peroxisome movements

To quantify peroxisome movements, time lapse images of *A. nidulans* strains expressing mCherry-Pts1 with or without overexpressed PxdA(Δ1-500)-TagGFP were collected at 500 ms intervals for 1 minute total. A line was drawn perpendicular to the hyphae 10 $\mu$m from the hyphal tip. The number of peroxisomes that crossed the line over a 60 second movie was counted [12]. A Mann-Whitney test was performed to determine statistical significance.

#### S5. SIMULATING LINKER PROTEIN ENCOUNTERS

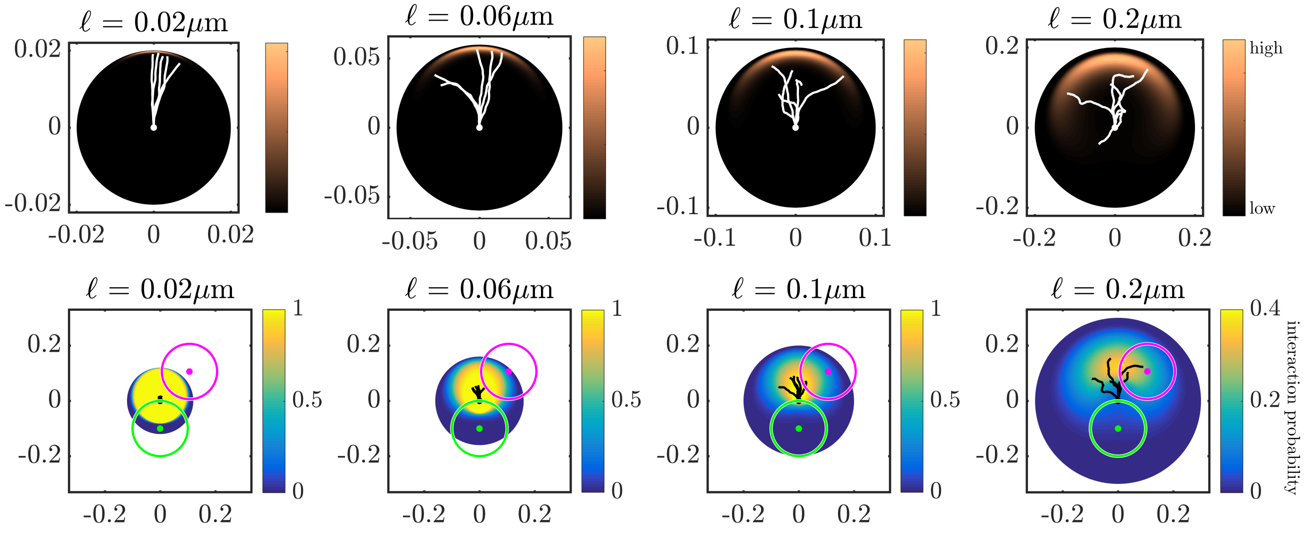


Figure S4: (Top) Spatial distribution for the free end of a wormlike chain with persistence length $\ell_p = 0.1\mu$m, where one end has its position fixed at the white dot and orientation fixed upwards. Color indicates the probability density of finding the linker tip at a given location. Sample configurations of linkers are shown with white lines. (Bottom) Probability of contact [$p(d,\beta)$] between a cargo organelle and a linker protein, shown as a function of cargo position relative to the linker attachment point (black dot). The carrier organelle location (green), example cargo location (magenta), and sample linker configurations (black lines) are shown. Both cargo and carrier have a radius of $0.1\mu$m.

In *A. nidulans* hyphae, the protein PxdA is a putative linker that connects hitchhiking peroxisomes to motor-driven early endosomes. Analysis of the makeup of PxdA reveals a coiled-coil region that is predicted to be about $\approx 90$nm in length[3]. We also note that the persistence length of coiled-coil proteins is reported to be in the range $30 - 170$nm [13–16]. The expected mechanical properties of these linker proteins thus fall in the category of semiflexible polymer chains, and we therefore model a typical linker protein as a "worm-like" chain[17] (WLC) whose base is fixed on the surface of the carrier organelle.

Using analytically derived Greens' functions, we first obtain the spatial probability distribution $G(r,\theta,\phi;\hat{z})$ of the linker end point, assuming the initial orientation ($\hat{z}$) points radially outward from the carrier [18]. Since the distribution is azimuthally symmetric, we can represent the distribution in terms of $r$ and $\theta$ as $G'(r,\theta;\hat{z}) = 2\pi G(r,\theta,\phi;\hat{z})$. Note that this representation preserves the normalization $\int_0^{2\pi}\int_{-1}^{1}\int_0^{\ell} r^2 G(r,\theta,\phi;\hat{z})\,dr d(\cos\theta) d\phi = 1$. Fig. S4 (top) shows the end distribution $G'(r,\theta;\hat{z})$ for a WLC with persistence length $l_p = 0.1\mu$m, and varying chain lengths ($\ell$).

To determine the probability of contact between the linker and a hitchhiker located at any given position, we calculate the overlap between the equilibrium spatial distribution of the WLC tip and the volume occupied by the hitchhiking cargo. In the reference frame of the fixed linker end, we assume a sphere of radius $r_p$ is placed at polar coordinates $(d,\beta,0)$, where $d$ is the distance from the linker attachment point and $\beta$ is the angle of the hitchhiker position relative to the initial linker orientation. The integral of the linker end distribution over the volume occupied by the sphere then gives the probability of contact for that particular position of the hitchhiking cargo.

Assuming the tip of the WLC is located at $(r,\theta,\phi)$, its distance from the center of a hitchhiker located at $(d, \beta, 0)$ is given by

$$R = \sqrt{r^2 + d^2 - 2rd(\sin\theta\cos\phi\sin\beta + \cos\theta\cos\beta)}$$

For the point $(r, \theta, \phi)$ to be in the interior of the hitchhiker, the distance $R$ should be less than the hitchhiker radius, or $R < r_p$. Comparing these expressions gives us a range of possible values of $\phi$ for which the point $(r, \theta, \phi)$ lies inside the hitchhiker. The range of $\phi$ values satisfies

$$\cos\phi \geq \frac{r^2 + d^2 - r_p^2}{2rd\sin\theta\sin\beta} - \cot\theta\cot\beta.$$

The total amount of azimuthal overlap can be obtained from the values of $\phi$ that satisfy the above inequality. Denoting this overlap by $\Delta\Phi(r, \theta; d, \beta)$, we can write

$$\Delta\Phi(r, \theta; \phi, \beta) = 2\cos^{-1}\left(\frac{r^2 + d^2 - r_p^2}{2rd\sin\theta\sin\beta} - \cot\theta\cot\beta\right).$$

Note that $\Delta\Phi = 0$ if the linker tip cannot overlap with the hitchhiker, and $\Delta\Phi = 2\pi$ for complete overlap.

Using the azimuthal overlap, we can write the probability of interaction between the tip of the WLC and the hitchhiker as

$$p(d, \beta) = \int_{-1}^{1} d(\cos\theta) \int_0^{\ell} r^2 dr \frac{\Delta\Phi(r, \theta; d, \beta)}{2\pi} G'(r, \theta; \hat{z})$$

The probability $p(d, \beta)$ is tabulated with $d$ varying from $\approx 0$ to $(\ell + r_p)$, and $\beta$ varying from 0 to $\pi$. Fig. S4 (bottom) shows $p(d, \beta)$ for varying linker lengths. In our simulations, whenever the hitchhiker comes within a distance $(\ell + r_p)$ from the base of a linker, we calculate its position $(d, \beta)$ relative to that linker base and obtain the associated overlap probability $p(d, \beta)$ by interpolating the tabulated values. Successful contact is determined by sampling from this probability at each time-step.

#### S6. EXTRACTING THE RATE OF CONTACT BETWEEN CARRIER AND HITCHHIKER

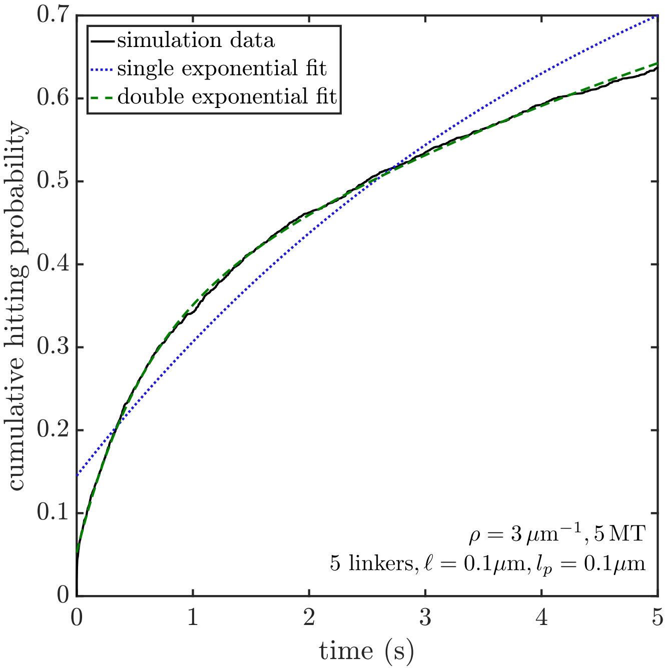


Figure S5: Cumulative contact probability (solid black line) vs time showing single (blue, dotted line) and double (green, dashed line) exponential fits.

The rate of contact between the carrier and the hitchhiker is extracted from the cumulative probability of contact vs time obtained from simulations. As described in the Methods section, 2500 trials of carrier passage in a tubular geometry are simulated with Brownian dynamics. For each trial, we record the probability of contact occurring within a given time-step. This probability density is integrated to obtain the cumulative probability distribution as a function of time, which is then fit to a double exponential function $F(f_1, \tau_1, f_2, \tau_2, t) = 1 - f_1 e^{-t/\tau_1} - f_2 e^{-t/\tau_2}$. The two weights $f_1, f_2$ need not sum to 1, because some trajectories start with the cargo already in contact with a carrier organelle or linker tip. The average time to contact is obtained as $\tau = f_1\tau_1 + f_2\tau_2$, and the overall rate of contact is defined by $k_{\text{hit}} = \tau^{-1}$. This approach effectively extrapolates the cumulative contact time distribution to longer times using a double-exponential fit. It allows the calculation of an effective rate even when that rate is quite small, without the need for the very long simulations that would be required for all trials to achieve contact. The more commonly used single exponential fitting function $[F(f, \tau, t) = 1 - f e^{-t/\tau}]$ provides a poor fit compared to the double exponential (Fig. S5). We note that the qualitative behavior of the rate of contact between a hitchhiker and carrier remains the same regardless of the choice of fitting function.

#### S7. CONTACT RATES FOR LARGER HITCHHIKERS

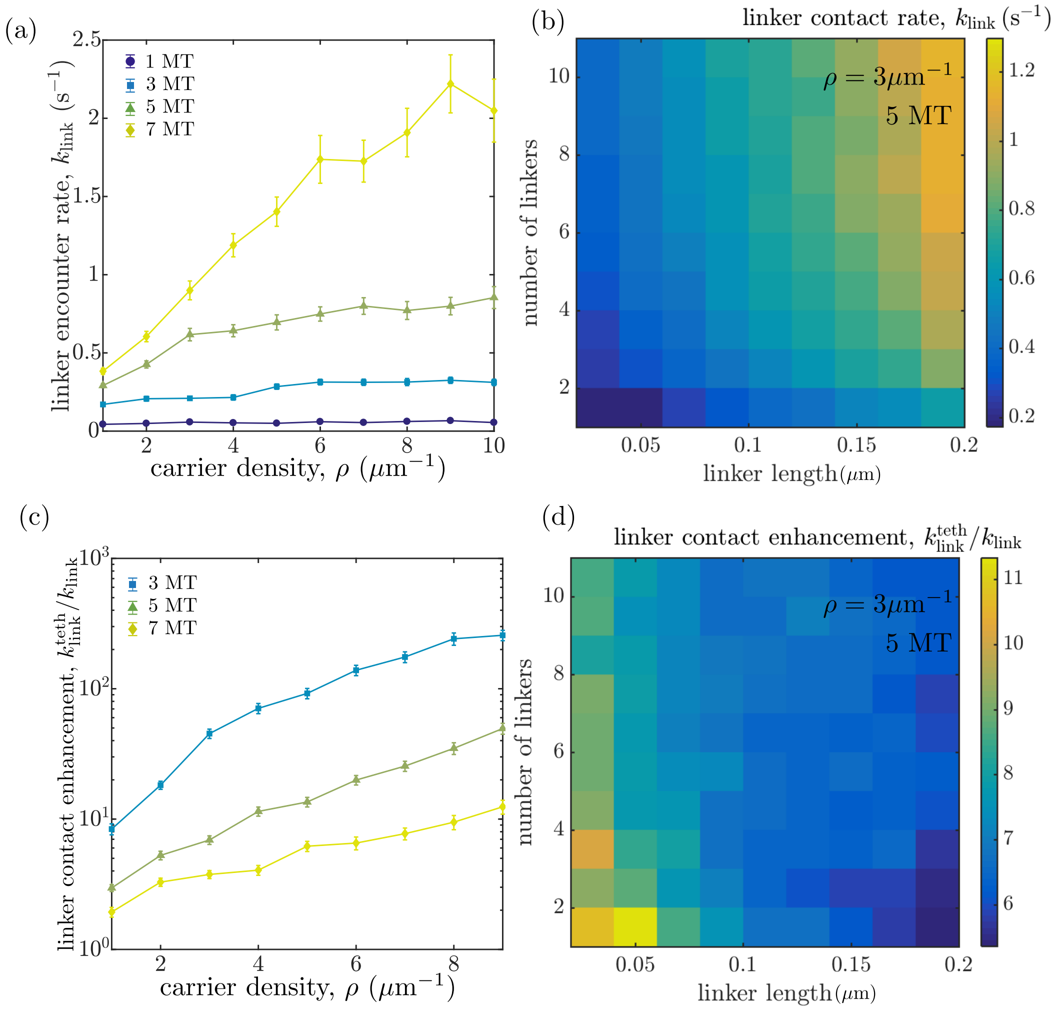


Figure S6: Rates of linker encounters for larger hitchhiking organelles ($r_p = 0.3\mu$m). (a) Rate at which hitchhiker encounters the first linker tip as a function of carrier density and the number of microtubules. (b) Effect of linker length and linker number per carrier on the encounter rate. (c) Enhancement of overall contact rate with a linker protein tip, due to tethering of cargo. (d) Ratio of encounter rates with linker protein tips, for a tethered versus diffusive cargo.

Peroxisomes are dynamic organelles that can vary in size ranging from 0.1$\mu$m-1$\mu$m depending on the cell type. Peroxisomes are also known to modify their interactions with other subcellular components resulting from morphological changes [19]. In this section, we explore how changing the size of the hitchhiking cargo affects the rate of encounter with passing carriers.

For the tubular geometry described in the main text, increasing the size of the hitchhiker enables it to access a larger cross-section of the intracellular space, making it more likely to come in contact with a carrier. Assuming the radius of the hitchhiker as 0.3$\mu$m (compared to 0.1$\mu$m used in the main text), we calculate the rate of encounter with the tip of a linker on a carrier organelle. We note that while the overall rate of encounter increases, the qualitative dependence on carrier density, linker length and number remains the same.

Exploring the effect of tethering large hitchhikers to microtubules, we note that the enhancement of the encounter rate due to tethering is somewhat diminished compared to smaller hitchhikers. However, the overall enhancement still remains substantial and can be up to 8-fold for carrier transport parameters discussed in the main text (compared to $\approx$12-fold for hitchhikers of radius $0.1\mu$m).
